# ‘Keeping the kids at home’ can limit the persistence of contagious pathogens in social animals

**DOI:** 10.1101/2020.04.11.036806

**Authors:** L. Marescot, M. Franz, S. Benhaiem, H. Hofer, M.L. East, S. Kramer-Schadt

**Author notes:** equal contribution (L. Marescot, M. Franz, S. Benhaiem). equal contribution (M.L. East, S. Kramer-Schadt). Bundesinstitut für Risikoforschung (BfR), Diedersdorfer Weg 1, 12277 Berlin, Germany (M. Franz).

## Abstract

In social species where offspring are reared together in communal burrows or similar structures, young animals typically do not engage in between-group contact during their development – a behavioural trait we call ‘offspring with restricted between-group contact’ (ORC). The impact of this trait on the persistence of contagious pathogens that generate lifelong immunity in their hosts is currently unclear. We hypothesize that in populations with ORC, the formation, in groups, of a ‘protective barrier’ of only recovered adults, prevents the transmission of this type of pathogens to the new susceptible hosts (i.e. young animals), thereby increasing the probability of epidemic fade-out. We implement a spatially implicit individual-based Susceptible-Infected-Recovered (SIR) model for a large range of host and pathogen traits and show that the epidemic fade-out probability is consistently higher in populations with ORC, especially when disease spread is fast (high basic reproduction number R_0_). We also show that ORC can counteract the cost of group-living in terms of disease risk to a greater extent than variation in other traits. We discuss our findings in relation to herd immunity and outline how they could be used to implement efficient management measures such as vaccinations.

## INTRODUCTION

Pathogens of infectious diseases that spread through direct contact or physical proximity can imperil species that live in dense populations or large social groups, including humans, livestock or wildlife (Altizer et al. 2003, e.g. the COVID-19 pandemic in humans: Anderson et al. 2020). Understanding the patterns of transmission and the mechanisms of persistence of such diseases in group-living species is thus of key relevance for human public health, economy or wildlife conservation. The structure of social contact networks in group-living species may critically influence the spread and persistence of contagious diseases (Kappeler et al. 2015). For example, if a social network contains a few central individuals who are exceptionally well connected to others, then these individuals can act as super-spreaders and their presence can strongly increase the likelihood and extent of an epidemic as well as the persistence of pathogens (Lloyd-Smith et al. 2005).

A social network can further be characterized by its modularity, that is, its degree of division into modules such as social groups (e.g. clans, prides or troops) or subgroups (e.g. age classes in the case of eusocial insects, Stroeymeyt et al. 2018, also see Griffin and Nunn 2012, Sah et al. 2017). Generally, a social network is considered to be highly modular, or to have a high ‘community structure’ (White et al. 2017) when individuals within one module are more connected to each other than they are to individuals in other modules (Newman 2006). It has been hypothesized that a highly modular social network can generally reduce disease risk in group-living animals (Griffin and Nunn 2012, Nunn et al. 2015). Yet this view has been recently challenged by a study which found that the beneficial effects of modularity on disease fade-out are restricted to extreme cases of exceptionally high modularity, which are rarely observed in natural systems (Sah et al. 2017).

Here we investigate how highly modular social networks characterized by age-dependent between-group contact affect pathogen spread and persistence. We focus specifically on networks where young animals have no contact with members of other groups, and term this life history trait ‘offspring with restricted between-group contact’ (ORC).

In contrast to most ungulate species for instance, in which precocial young follow their mother shortly after birth (Lent 1974), several group-living species rear their offspring in communal burrows, nurseries, crèches or family groups within group territories or home ranges for extended periods of time, during which time young rarely if ever contact individuals of other groups. This includes for instance spotted hyenas *Crocuta crocuta* (Kruuk 1972), African lions *Panthera leo* (Schaller 1972) or European rabbits *Oryctolagus cuniculus* (Daly 1981).

Restricting contact of young animals with individuals of other groups may specifically limit the persistence of pathogens that induce lifelong immunity following infection. Such pathogens, which include many viruses such as measles and canine distemper virus in the genus *Morbilivirus*, mumps in the genus *Rubulavirus* or rubella in the genus *Rubivirus* (Morris et al. 2015) rely on the birth of new susceptible hosts and their transmission to young individuals to persist in a host population (Lloyd-Smith et al. 2005).

An ORC trait could affect pathogen persistence via at least two, mutually non-exclusive, mechanisms. First, as a rather trivial effect, by restricting contact of young animals with members of other groups, ORC could directly reduce the probability of disease spread and hence also the basic reproduction number R_0_ of a pathogen, which is the expected number of secondary infections that a single infected host causes in a population of susceptible individuals (McCallum et al. 2001). Second, ORC could increase the chance of epidemic fade-out by depriving the pathogen of access to susceptible individuals during the late stage of an epidemic. Indeed, when most adults in the population have become immune (and thus herd immunity (see e.g. Anderson and May 1985) is high, the chance of pathogen spread to mostly susceptible young animals would be substantially curtailed. In other words, ORC may increase the efficiency of a protective barrier in groups composed of only recovered adults ─ which emerges naturally when social hosts are infected with Susceptible - Infected - Recovered (SIR) types of pathogens ─ thereby increasing the probability of epidemic fade-out.

Here we used an individual-based model to investigate for the first time this second mechanism and test the prediction that the probability of epidemic fade-out increases in populations (i.e. networks) of social species with an ORC trait. For this purpose, we developed an SIR model to describe pathogen transmission and host population dynamics. In order to assess how consistently the probability of epidemic fade-out increased due to ORC, we systematically varied parameters likely to influence pathogen transmission dynamics: (1) host birth rate, (2) host death rate (which occurs independently of infection), (3) contact rates between members of different groups, (4) infection rate at contact, (5) individual infection length, (6) virulence (i.e. the increase in host death rate due to infection), and (7) whether transmission is frequency or density-dependent. For each specific combination of parameters, we investigated two scenarios. First, we set up a baseline scenario in which rates of between-group contact were identical for individuals of all ages. Second, we investigated an ORC scenario in which individuals below a given age-threshold could not contact members of other groups, i.e. their rates of between-group contact were set to zero. This is simulating the legal requirements of confinement put into action by authorities to limit the spread of pandemics, e.g. during the COVID-19 pandemic; the closure of schools and prevention of contact between young and elderly people, including within households (e.g. Anderson et al. 2020, Kelvin and Halperin 2020). Because ORC decreases R_0_, we adjusted the infection rate in the baseline scenario such that the value of R_0_ was always identical in both scenarios to insure that the emergence of a more efficient immunity barrier are directly related to the ORC trait, and not to its mediated effect on R_0_. We discuss our results in relation to behavioural mechanisms yielding herd immunity as well as the likely outcomes of altering social network modularity through management measures.

## METHODS

### Model description

The model description followed the ODD (Overview, Design concepts, Details) protocol for describing individual-and agent-based models (Grimm et al. 2006, Grimm et al. 2010). The model was implemented in NetLogo (v.5.2.0), a free software platform for agent-based simulations as an integrated modeling environment, and is available on GitHub (https://github.com/LucileMarescot/RestrictingOffspringBetweenGroupContact_IBM). We used R v. 3.5.3. (R Core Team 2019) and the packages *ggplot2* (Wickham 2016), *tidyverse* (Wickham et al. 2019a), *dplyr* (Wickham et al. 2019b), *patchwork* (Pedersen 2019), *showtext* (Qiu 2020), *here* (Müller 2017) and *ggtext* (Wilke 2020) to produce the figures.

#### Purpose

The purpose of our model was to study how disease dynamics in group-living species are affected by an ‘offspring with restricted between-group contact’ (ORC) trait, which occurs when offspring are kept in subunits or ‘modules’ such as communal dens and burrows, crèches or family groups within their social groups, until they reach the age at which they can start engaging in between-group interactions. We were particularly interested in whether, and to what extent, such age-dependent patterns of between-group interactions consistently increased the probability of epidemic fade-out across a large range of host and pathogen life history traits. We used life history parameter estimates from previous research findings on a free-ranging population of a highly social carnivore, the spotted hyena, in the Serengeti National Park in Tanzania. This population presents age-dependent patterns of between-group interactions: offspring are stationed at (or remain in the vicinity of) a communal den inside the territory of their social group (clan) for ca. the first 12 months of life. Juveniles then start roaming more widely within the clan territory, and to accompany their mothers on regular long-distance foraging trips outside the clan territory to areas containing large aggregations of migratory herds (Hofer and East 1993, Hofer et al. 2016). Spotted hyenas in the Serengeti National Park give birth throughout the year (East et al. 2003, Hofer and East 2003) hence communal dens typically contain susceptible young for much of the year.

Females remain in their natal clan whereas most males disperse when approximately four years old (East and Hofer 2001). However, we did not investigate sex differences in our model as they were not relevant to our question. We used the parameters survival probability, age at first reproduction and the probability of reproducing (from Marescot et al. 2018 and references therein). Please note that we did not develop a ‘hyena model’ *per se*, rather a realistic host-pathogen model to test our theory with (Railsback et al. 2020). All model parameters can easily be adapted to yield generalized results, or to tailor the model to more species-specific life histories.

#### Entities, state variables, and scales

Individuals and modules were regarded as entities. Each individual was characterized by three state variables, which were updated on a weekly basis: age, epidemiological status (susceptible S, infected I or recovered R, i.e. immune) and the time spent infected. Modules represented social groups, which consisted of a certain number of individuals, which could change over time due to processes such as mortality and dispersal.

#### Processing and scheduling

The model included two main processes: host population dynamics and disease dynamics. At each time step, the following sequence of processes occurred: (1) pathogen transmission, (2) death, (3) reproduction, (4) dispersal and (5) ageing of host individuals. We considered two classic forms of pathogen transmission (parameter *Transmission* in Table 1): density-dependent transmission, where the spread of infections depends on the size of the groups, and frequency-dependent transmission, where spread depends on contact frequency and not group size. Abbreviated model parameters (in italics below) are defined in Table 1, where their simulation values are also provided.

**Table 1:** Host and pathogen-specific parameters in the individual-based model. The numbers in square brackets represent the values explored for each parameter that was varied in order to assess the sensitivity of epidemic fade-out probabilities to changes in host and pathogen traits. The unit of time was week and state variables were updated on a weekly basis.

| Parameter | Description | Simulation values [values] |
| --- | --- | --- |
| **Host** |  |  |
| $K$ | Carrying capacity of the population | 1000 |
| $Group.size$ | Carrying capacity of the group; i.e. group size threshold | [10, 50, 100], leading to [100, 20, 10] numbers of groups initialized |
| $Age-contact$ | Age at first between-group contact (baseline: 0 week, ORC: 52 weeks) | [0, 52] |
| $Age-repro$ | Age at first reproduction (adults only) | 104 |
| $\phi$ | Survival probability | [0.991, 0.995, 0.998] |
| $f_{age}$ | Probability of reproducing, resulting in a fertility of 1-3 offspring p.a. | [0.01, 0.025, 0.05] |
| $\pi$ | Contact ratio: ratio between the rate of between-group contact and the rate of within-group contact | [* , 0.1, 0.5, 1] |
| **Pathogen** |  |  |
| $Inf$ | Infectivity or ability of the pathogen to infect the host; adjusted to obtain similar $R_0$ values for baseline and ORC scenarios | [1, 2, 5, 10, 20] |
| $\alpha$ | Virulence (probability of the host to die from an infection; case fatality) | [0, 0.05, 0.1] |
| $IL$ | Infection length in weeks | [10, 20, 30] |
| $Transmission$ | Frequency-dependent vs. density-dependent transmission mode | Frequency-dependent, density-dependent |

\* We did not model the case of $\pi = 0$ , as this would imply that the pathogen is contained within one group and always dies out.

In the first process (pathogen transmission), the force of infection was influenced by i) the age at first between-group contact, *Age-contact*, ii) the number (for density-dependent transmission) or proportion (for frequency-dependent transmission) of infected individuals and iii) the contact ratio, *π*, which calculates the ratio of between-group to within-group contact rate. This ratio defines how much two individuals from different groups interact compared to two individuals of the same group. That is, if *π* = 1, all individuals interacted equally, independently of their social group affiliation (low modularity). Conversely, if *π* = 0, social interactions occurred exclusively within groups. In the ***baseline scenario*** we assume equal contact between all age classes, with contact rates within and between groups defined by *π*. In the ***ORC scenario***, i.e. the case when young individuals started to engage in between-group contact only from a given age (here *Age-contact* = 52 weeks), modularity in the network was high. For simplicity, hereafter we use ‘juveniles’ to refer to individuals younger than 52 weeks (i.e. individuals that have no contact with members of other groups in the model with ORC) and ‘adults’ to refer to individuals older than 52 weeks (i.e. individuals which have contact with members of other groups in the model with ORC). Finally, pathogen traits such as virulence, *α*, infectivity, *Inf*, and maximum infection length, *IL*, also affected the transmission pattern. The social network was restructured following pathogen transmission, due to the processes described below.

The second process (death) included two forms of mortality: the host intrinsic mortality (1-*ϕ*, with *ϕ* being the weekly survival probability) and the host mortality induced by the pathogen (virulence *α*). Individuals who survived over the maximum infection length (*IL*) gained immunity for the rest of their lives.

The third process (reproduction) was regulated by the carrying capacity of the host *K*. Births occurred only in groups that were below a local breeding capacity, which was defined by the maximum group size. The probability that an individual at a reproductive age of 2 years and more (*Age-repro ≥* 104 weeks) gave birth to one offspring was given by *f*_*age*._ and corresponded to approximately 1 – 3 offspring per year.

The fourth process (dispersal) described individuals which left their natal group as they became reproductively mature.

In the last process (ageing), we updated the individual’s state variable *age* as well as the time-span-infected counter.

#### Initialization

The initialization of the age of individuals was determined by a random exponential distribution with a mean set at the age at first reproduction (*Age-repro =* 104 weeks, Table 1). All groups were set with a certain size and all individuals were set as susceptible, except one who was set as infected. Pathogen invasion started stochastically with this one infected individual chosen randomly in the initial population. The total population was at a carrying capacity of 1000 individuals at the beginning of the simulations to make simulation runs comparable, but the number of groups was given depending on the initialized number of individuals per group (see also Table 1).

#### Input data

Except for life history parameter estimates from previous studies on spotted hyenas (from Marescot et al. 2018 and references therein, as mentioned above), this theoretical model did not use any empirical input.

#### Submodels

##### (1) Pathogen transmission

For individuals who reached their age of first between-group contact (ORC: *Age-contact* = 52 weeks, baseline scenario: *Age-contact* = 0 week, Table 1), infection could occur both within (termed ‘inside’) and between (termed ‘other’) groups. Both sources of infection were considered in the calculation of the force of infection *β*_*i,age ≥ Age-contact*_ of an individual *i* for frequency-dependent transmission:

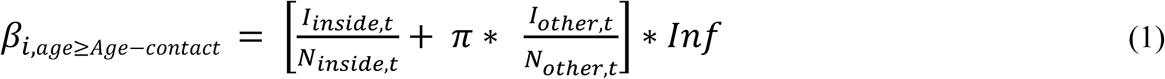

And for density-dependent transmission:

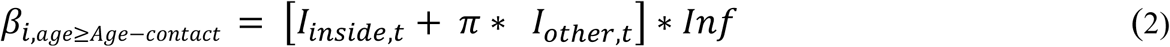

In both equations the first term in the brackets represented the contact rate between a susceptible individual from a group with infected individuals from the same group at a time step *t* (*I*_*inside,t*_). The second term in the brackets represented the contact rate between a susceptible individual from a group with infected individuals from other groups (*I*_*other,t*_). This second term depended on the contact ratio *π*, hence accounted for contact rates between individuals from different groups (see Table 1). Both terms were then multiplied by the infectivity of the pathogen *Inf*, describing the probability that a susceptible host became infected after exposure to the pathogen (see Table 1). In (1), *N*_*inside,t*_ and *N*_*other,t*_ represent the densities within a given group and in other groups, respectively. For a susceptible individual *i*, the probability of becoming infected *P*_*i*_ was given by:

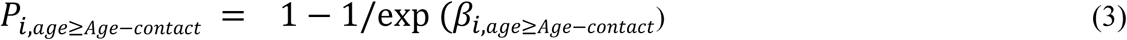

For a susceptible individual *i* of age < *Age-contact* contact, the probability of becoming infected depended on the infectivity of the pathogen and the risk of exposure to an infected individual.

For frequency dependent transmission, the force of infection *β*_*i,age ≤Age-contact*_ was thus:

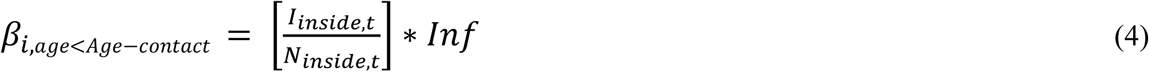

For density-dependent transmission:

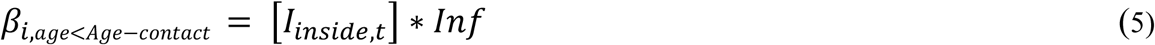

Each susceptible juvenile *i* therefore had a probability of becoming infected *P*_*i*_ given by:

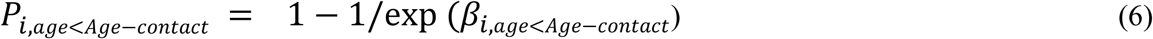

When an individual was infected, its epidemiological status changed from susceptible to infected and the counter for the time spent infected was set to one (for the first week of infection), and then increased in each subsequent week of infection. If the individual had survived the entire duration of the infection period (*IL*), its epidemiological status changed from infected to recovered. Pathogen clearance occurred when individuals were free of infection and still alive at the end of the infection period *IL*, i.e. when they entered the recovered state. We did not consider a constant pathogen spillover from reservoirs nor a latency period after exposure to infection.

##### (2) Death

At each time step, all individuals died with a probability 1-*ϕ*, with *ϕ* being the survival probability, which captured causes of intrinsic mortality. In addition, infected individuals died with probability *α*, i.e. the virulence of the pathogen.

##### (3) Reproduction

We assumed density-dependent reproduction. Reproduction only occurred if the group size was below the carrying capacity of groups, i.e. the *Group*.*size* threshold (Table 1). If reproduction occurred, then every week, each adult which was older than *Age-repro* gave birth to a single offspring with a probability *f*_age_. All individuals were born as susceptible, independently of the infection status of their parent; hence, we did not consider vertical transmission or immunity due to the transmission of (maternal) antibodies to offspring (through e.g. milk) for a short period after birth.

##### (4) Dispersal

We modeled primary dispersal, i.e. all individuals left their natal groups when they reached their reproductive maturity *Age-repro* and moved to a randomly selected group (which included extinct groups).

##### (5) Ageing

The process of ageing updated the individual’s state variable age as well as the time-span-infected counter, which was the time in week(s) since an individual got infected.

### Model analyses

We aimed to assess whether an ORC trait increased the probability of epidemic fade-out consistently across a range of host and pathogen life history traits, while keeping the basic reproduction number of the pathogen, R_0_, identical in both scenarios (baseline and ORC scenario). To do so, we designed a multifactorial simulation experiment. We varied the simulation values of the parameters specific to the host life history: survival *ϕ*, fertility of reproductive individuals *f*_*age*_, as well as group-size (*Group*.*size*) and the contact ratio *π* (Table 1). We also considered variation in the simulation values for the parameters characterizing the pathogen: virulence, α, infection length, *IL*, and the basic reproduction number R_0_ that we calculated analytically (see S4 and S5, Supplementary material). We adjusted the infectivity rate *Inf* between the ORC and the baseline scenario (i.e. no age-dependent between-group contact) so that both types of networks had similar R_0_, all else being equal. We considered 2916 combinations of parameters. We ran each simulation over 10 years and recorded whether the epidemic persisted or faded out. We repeated each parameter combination 30 times to examine the probability of epidemic fade-out. We plotted the average epidemic fade-out probability for ORC and baseline scenarios, across all parameter combinations and for each value of R_0_.

To determine if the age at first between-group contact was the trait whose variation had the most important impact on epidemic fade-out probability, we compared epidemic fade-out probabilities calculated for the extreme values of each parameter, holding all other parameters equal. To avoid naïve comparisons of fade-out probabilities owing to the limited number of receptions of our simulations that could generate substantial bias in the calculated probabilities, we filtered the results by running a Fisher’s exact test for each paired scenario. We tested if differences in epidemic fade-out probabilities were significant and plotted the distributions of differences between epidemic fade-out probabilities only for the paired scenarios with a *p*-value < 0.05.

Finally, we ran a regression tree with the R package *rpart* (Therneau and Atkinson 2018), using the ANOVA method, to partition the difference in epidemic fade-out probabilities between the ORC and the baseline scenario. We used this approach to detect under which specific conditions (i.e. trait values) ORC showed the greatest effect – provided it did so (Breiman et al. 1984). The method consisted in splitting the parameter space into binary groups, which differed in terms of difference in epidemic fade-out probabilities with a minimum deviance of the standard deviation of the epidemic fade-out in those groups.

## RESULTS

### Effects of restricting offspring between-group contact on epidemic fade-out probability

As theory predicts, epidemics faded-out at a R_0_ < 1, and did so regardless of the type of between-group interactions. The probability of epidemic fade-out was consistently higher in hosts with an ORC trait than in the baseline scenario, and this difference increased with increasing R_0_ (figure 1).

**Figure 1:**
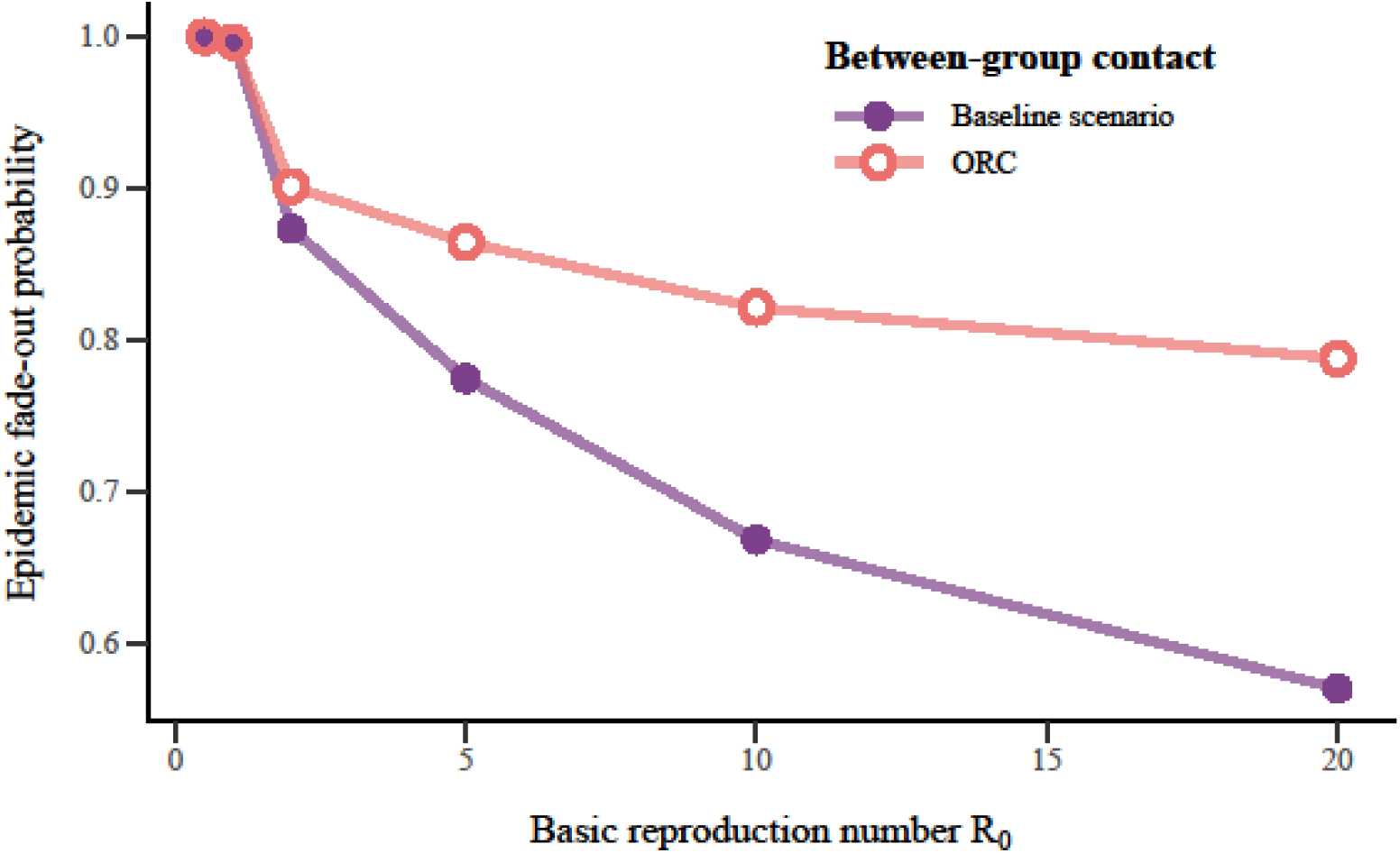
Relationship between the probability of epidemic fade-out and the basic reproduction number R_0_ in populations describing the two types of social interactions: age-dependent (offspring with restricted between-group contact (ORC), pink) and age-independent (baseline scenario, purple) between-group contacts, for all possible combinations of host and pathogen traits.

Age at first between-group contact was the parameter with the most important influence on epidemic fade-out probability. The difference in epidemic fade-out probabilities between a host with ORC (age at first between-group contact equal to 52 weeks) and a host without ORC (‘baseline scenario’) was indeed greater than the difference in epidemic fade-out probabilities between two most extreme values of any other parameter (figure 2, panel ‘Age at first between-group contact’).

**Figure 2:**
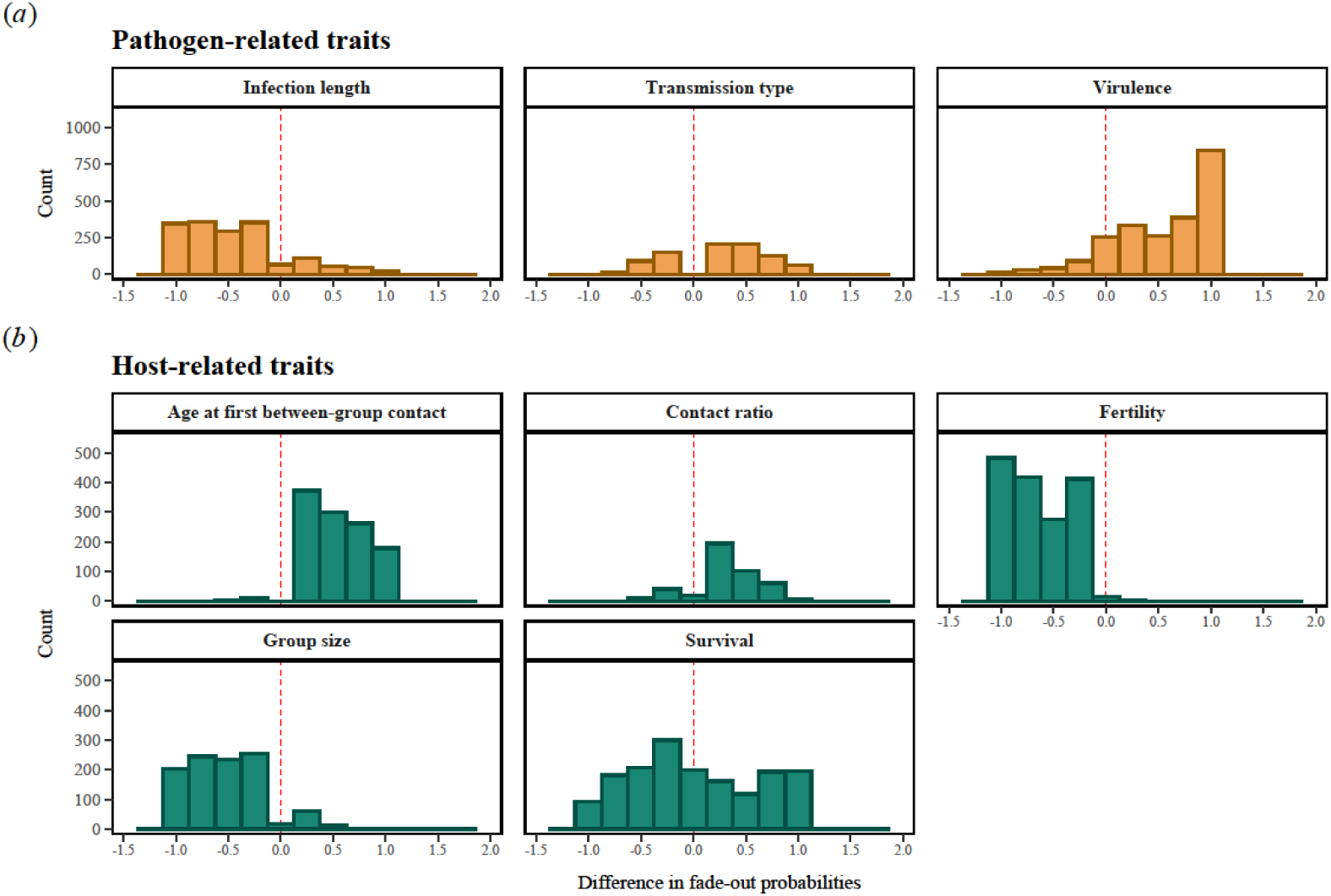
Number of simulations (count) in relation to the difference in epidemic fade-out probabilities between two extreme values of each parameter (for (*a*) pathogen-related traits (orange) and (*b*) host-related traits (blue-green)), all else being equal. The extreme values of each parameter were as follows: (*a*) infection length (IL = 30, IL = 10), transmission type (frequency vs density) and virulence (*α* = 0.05, *α* = 0); (*b*) age at first between-group contact (52 weeks for ORC, 0 for baseline scenario), contact ratio (π = 1 and π = 0.1), fertility (*f*_*age*_ = 0.05, *f*_*age*_ *=* 0.01), group size (100, 10) and survival (*ϕ =* 0.998, *ϕ =* 0.9910). A positive difference (on the right side of the horizontal dotted line) indicates that the scenario with the highest value for a given parameter results in a higher epidemic fade-out probability than the scenario with the lowest value for that parameter. This figure shows that the difference in fade-out probabilities is almost always (highly) positive only for the parameter age at first between-group contact.

The positive effects of ORC for the host population in terms of epidemic fade-out probability were generally more important at high values of R_0_ and for host species with a high fertility rate (figure 3). Given this set of conditions, benefits were then particularly elevated (1) for pathogens with a short period of infection and density dependent transmission (0.6 and 0.51, figure 3), (2) in social networks composed of small groups affected by pathogens with a long period of infection and low virulence (0.47, figure 3) and (3) networks with low modularity, frequency dependent transmission and a short period of infection (0.37, figure 3).

**Figure 3:**
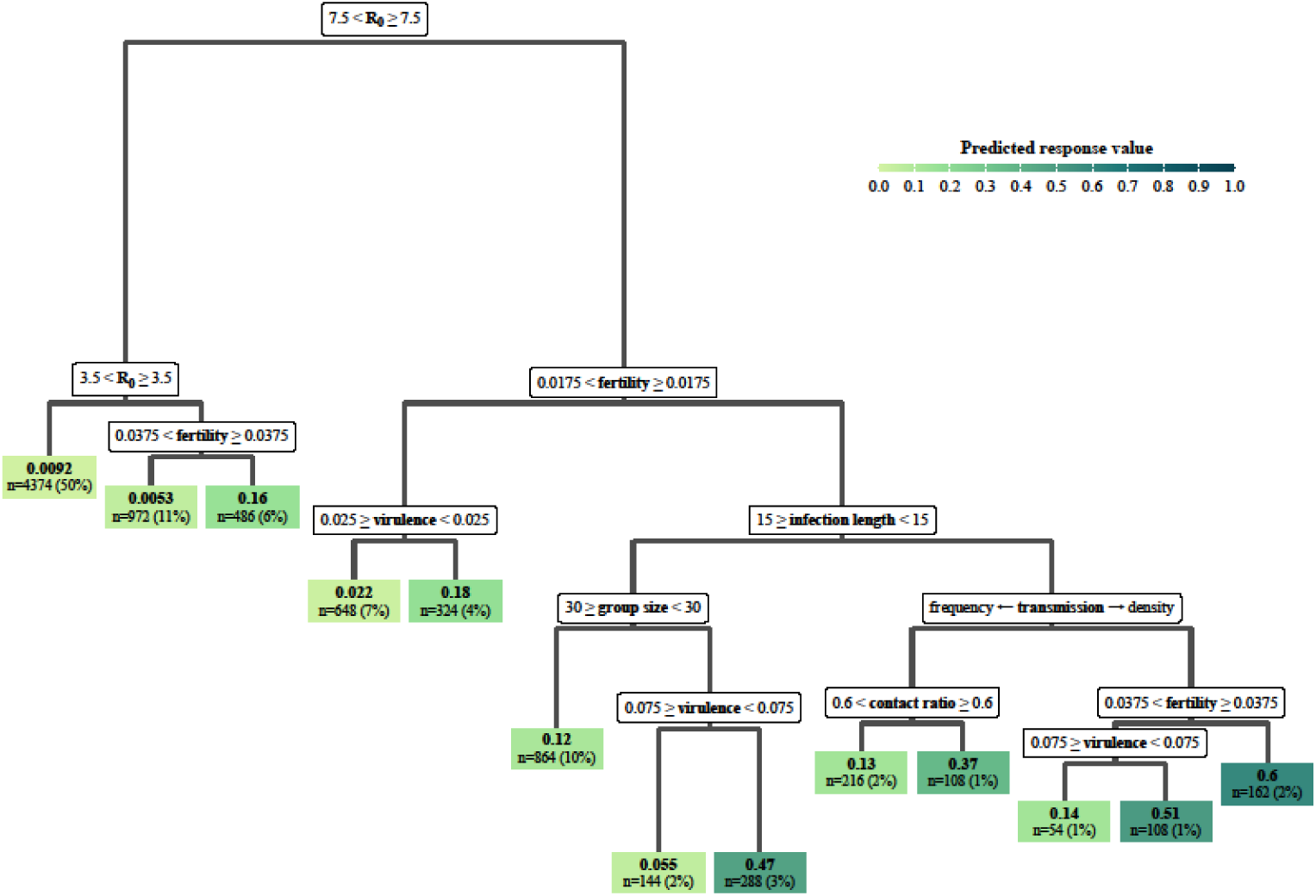
Regression tree of the differences in the epidemic fade-out probability between host populations whose ‘offspring had restricted between-group contact’ (ORC) vs those who did not, across all simulated parameters. The method consisted in splitting the parameter space into binary groups, which differed in terms of difference in fade-out probabilities successively, using the ANOVA method. At each internal node, the label indicates the parameter considered. For instance, the first split at the top of the tree results in two main branches: the left one corresponds to R_0_ < 7.5 and the right branch corresponds to R_0_ ≥ 7.5. The values at the terminal nodes indicate differences in epidemic fade-out probabilities.

### SIR dynamics

We found important differences in how the number of individuals in the susceptible, infected and recovered epidemiological states changed over the first 10 years (520 weeks), between the ORC and the baseline scenario (figure 4). When looking at a single epidemic event in a host population with ORC, we found that a new pool of susceptible juveniles (free from the disease) emerged in the population a few weeks after initialization to form a peak, followed by an increase in the number of susceptible adults (figure 4 (*a*)). The epidemic faded out from the host population approximately 50 weeks after the initialization, both in terms of the number of infected juveniles and infected adults (figure 4(*b*)). This corresponds approximately to the age at which juveniles in the first cohort became adults and thus started contacting (potentially infected) members of other groups (as *Age-contact* = 52 weeks, see Table 1). The host population rapidly reached a stage in which all adults became recovered (approximately 27 weeks in figure 4 (*c*)). The last infected individuals were juveniles, and those surviving the infection became recovered adults (figure 4 (*c*)). Consequently, a few weeks after pathogen extinction and population turnover, the new pool of only susceptible juveniles emerged in the population (figure 4 (*a*)). The number of recovered adults declined over time as they were not fuelled anymore by infected juveniles who turned into recovered adults (figure 4 (*c*)).

**Figure 4:**
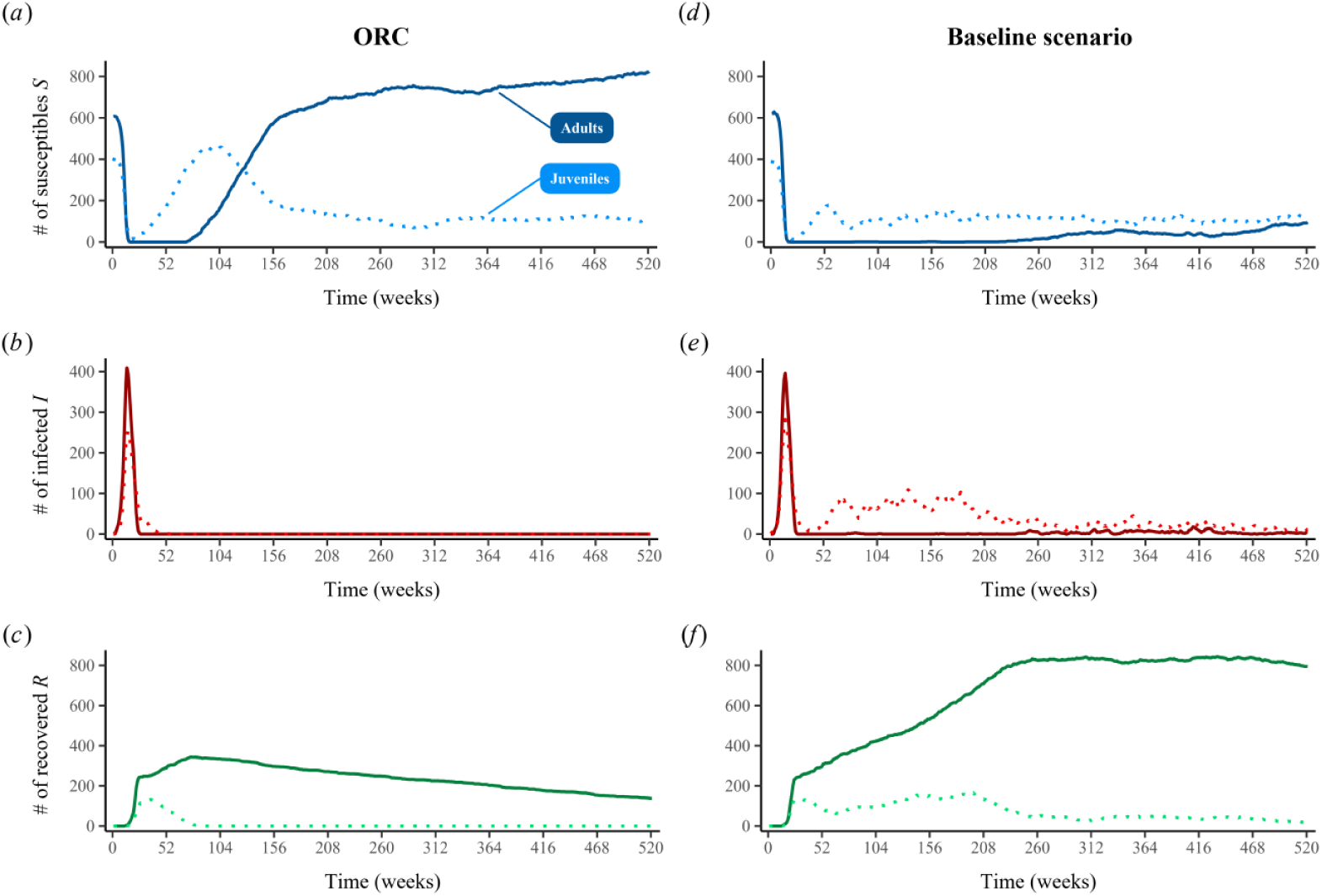
SIR dynamics during the first 10 years (520 weeks) of pathogen invasion in a host population with ORC (left) and in the baseline scenario (right): (*a*) and (*d*) number of susceptible juveniles (dotted blue lines) and adults (full blue lines); (*b*) and (*e*) number of infected juveniles (dotted red lines) and adults (full red lines) and (*c*) and (*f*): number of recovered juveniles (dotted green lines) and adults (full green lines). In both scenarios, we considered 20 groups of 50 individuals with a survival probability *ϕ* equal to 0.9, a probability of reproduction *f*_*age*_ equal to 0.05, resulting in an average fecundity of 2.5 offspring per year and a within-group contact 10 times higher than the between-group contact (π = 0.1). Regarding the value of pathogens traits, we assumed a virulence *α* of 0.1, an infection period *IL* of 10 weeks, a R_0_ of 10 and frequency-dependent transmission mode. This figure shows that the epidemic fades out very quickly in the population with ORC whereas the population in the baseline scenario faces frequent reinfections.

On the other hand, in the baseline scenario, when looking at the same epidemic event, we found that the host population was never composed of a large number of susceptible individuals (figure 4 (*d*))) and that the disease persisted for at least 10 years after the peak of infected juveniles and adults (figure 4 (*e*)). Despite a high number of recovered adults, the pathogen was maintained in the host population at a prevalence of approximately 2 % (figure 4 (*e*)). Although a large portion of adults was recovered and did not contribute to disease transmission (figure 4 (*f*)), the disease was maintained in the population as a result of the contact between infected and susceptible juveniles from different groups.

### Emergence of the protective barrier in groups

In the host population with ORC, during the first 200 weeks after pathogen invasion, the proportion of groups that did not contribute to the between-group transmission increased over time (figure 5 (*a*), (*b*)). These groups were ‘sealed’ because they were composed of adults who were all recovered and juveniles (susceptible, infected or recovered) who did not contact members of other groups. After approximately one year (100 weeks), all groups were sealed and composed of susceptible juveniles and recovered adults ─ a process which resulted in epidemic fade-out (figure 4 (*b*), figure 5 (*b*)).

**Figure 5:**
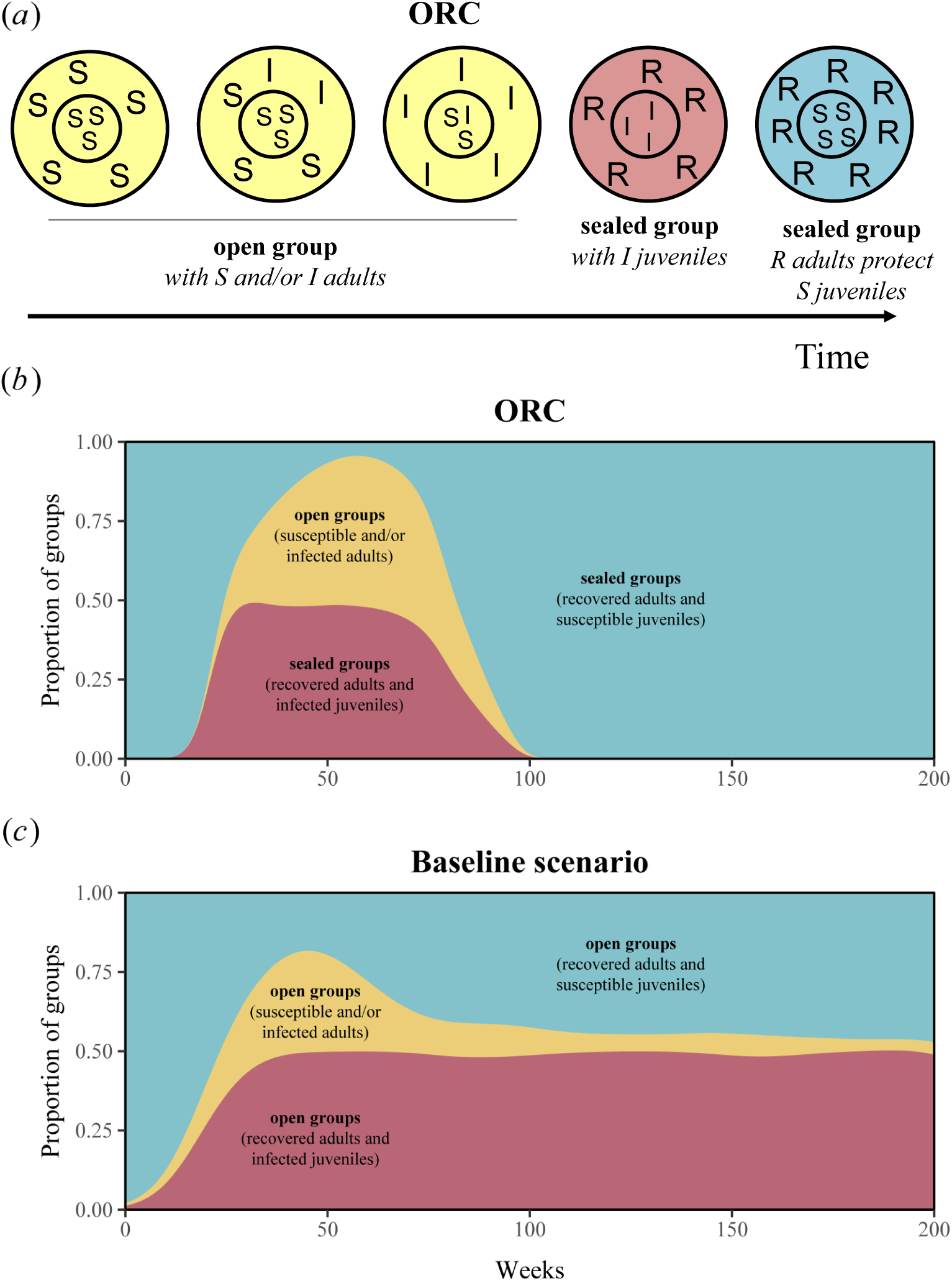
Emergence of protective barriers. Top/ Circles: (*a*) Schematic representation of temporal changes in the composition of groups in terms of susceptible (S), infected (I) and recovered (R) juveniles and adults, in a host population with ORC - where juveniles do not contact members of other groups. Large circles represent groups and encompass both juveniles (within the small circle, small capital letters) and adults (within the outside ring, large capital letters). Bottom/ Graph: Changes in the proportion of the different types of groups in the population during the course of the epidemic: (i) in yellow, groups in which all individuals are either susceptible or infected and can thus contribute to disease transmission (‘open groups’ in both the (*b*) ORC and (*c*) the baseline scenario), (ii) in pink, groups in which all adults are recovered and some juveniles are infected (‘sealed groups’ in (*b*) ORC and ‘open groups’ in (*c*) baseline scenario), and (iii) in blue, groups in which all adults are recovered adults and all juveniles are susceptible (‘sealed groups’ in (*b*) ORC and ‘open groups’ in (*c*) baseline scenario). We used the same trait values as for figure 4.

In contrast, in the baseline scenario, as susceptible juveniles could become infected when contacting members of other groups, all types of groups remained ‘open’ and contributed to disease transmission in the host population, either by having individuals getting infected, or by having individuals infecting other members of other groups. After 200 weeks, groups with susceptible juveniles and recovered adults (blue area in figure 5 (*c*)) and groups with infected juveniles and recovered adults (pink area in figure 5 (*c*))) were both roughly equally present in the host population.

## DISCUSSION

We demonstrated that across a wide range of host and pathogen traits, the probability of epidemic fade-out was consistently higher when juveniles had no contact with members of other groups (figure 1, figure 2), even after accounting for the negative effect of this trait on R_0_. Our results suggest that the formation of protective barriers composed of recovered adults around the new generation of susceptible juveniles (i.e. ‘sealed groups’) is the likely mechanism by which disease persistence is reduced in the population (figure 5 (*b*)). This phenomenon is expected to emerge during the late stage of an epidemic, when a high herd immunity, i.e. a high proportion of recovered individuals in the population (John and Samuel 2000), protects the new generation of susceptible juveniles from infection (Fine et al. 2011). The complete predominance of such sealed groups in the host population with ORC after several weeks was associated with epidemic fade-out (figure 5 (*b*), figure 4 (*b*)). We found that ORC had a particularly positive effect for the host population in terms of epidemic fade-out when R_0_ was elevated (e.g. equivalent to measles infection in humans, where R_0_ ∼ 18), and to hosts with high fertility rates and small group sizes (figure 3). This may be because the high production of offspring in small groups may further accelerate the formation of sealed groups and hence pathogen fade-out.

We investigated a specific form of network modularity, also called ‘community structure’ (Edmunds et al. 1997, Longini et al. 1982, Salathé and Jones 2010), where the structuring emerged from restricted contacts between groups. In accordance with the previous findings of Griffin and Nunn (2012) and of Nunn et al. (2015), who showed that when the structuring emerged from restricted contacts within groups, modularity slowed the spread of disease and reduced maximum prevalence, we found that highly modular social networks can reduce disease spread and persistence in group-living animals whose between-group contacts are restricted according to individual age. Sah et al. (2017) found that network modularity limited disease spread only when modularity was very high, i.e. when individuals within one module were more connected to each other than they were to individuals in other modules (Newman 2006). Here we showed that an ORC trait had a particularly positive effect for the host population in terms of epidemic fade-out for hosts with a high contact ratio, hence in networks of initially low modularity (figure 3). When the contact ratio was high, the pathogen could more easily spread between groups and was less likely to die out quickly; thus restricting offspring’s between-group contact could bring more benefits than when the contact ratio was low. These findings differ from those of Sah et al. (2017) possibly because we modelled host population dynamics. Thus in our model, changes in modularity stemming from the death of infected individuals or the birth of susceptible ones constantly altered the social network metrics during the epidemic ─ as modularity or centrality can only represent a snapshot of the community structure at one point in time.

In a disease management context, individuals with high contact rates to their group members are often targeted for vaccination to control the spread of infectious diseases (Carne et al. 2013). However, interventions focusing specifically on individuals bridging the modular network structure (i.e. with high between group contacts) may often be more efficient (Salathé and Jones 2010). Such ‘bridging’ individuals could be dispersing individuals in the case of rabies in foxes or raccoons (Reynolds et al. 2015), bovine tuberculosis (bTB) in badgers (Weber et al. 2013) or Classical Swine Fever (CSF) in wild boars (Scherer et al. 2020). Neglecting the potentially important influence of between-group interactions on disease spread, in combination with the disturbance of populations through persecution and hunting, may have been responsible for the failure of disease elimination programmes for bTB (Vial and Donnelly 2011) or CSF in Germany (Scherer et al. 2019). Our results also corroborate the so-called ‘network frailty concept’, showing the importance of immunization of highly connected individuals rather than random vaccinations as a management measure (Ferrari et al. 2006) and more generally, the need to consider the network structure of the underlying biological systems.

The ORC trait mimicked the case of hosts with clearly defined social units (e.g. groups) where offspring are reared together, thus have high contact rates with their own group members but no contact to members of other groups during their development. This definition suggests a spatial separation of the groups and clearly defined ‘nesting’ locations spaced at some distance, such as communal dens or burrows. In the Patagonian mara *Dolichotis patagonum* for instance, several pairs of adults cluster their young together in a crèche (Taber 1987, Ganslosser and Wehnelt 1997) (see Table 2 for other examples). Our results might also be applicable to cases where offspring are not stationary yet potentially protected from between-group contacts by other types of ‘barriers’, such as behavioural ones. This might include territorial space use of adults, e.g. troop formation in gorillas *Gorilla g. graueri* (Yamagiwa et al. 2003). In species with ‘mobile’ crèches and nurseries, such as common eider ducks *Somateria mollissima*, juveniles from different broods are raised together and guarded by adult females (McKinnon et al. 2006, Öst et al. 2007). It is likely that juveniles in such systems are more exposed to pathogens deposited by conspecifics in the environment (as some pathogens may survive several hours outside their hosts), as opposed to juveniles who are stationary in communal dens or burrows for a relatively long proportion of their lifespan. In contrast, colonial nesters, bird flocks, seabird colonies or the large herds of most ungulates such as zebras *Equus quagga*, wildebeest *Connochaetes*, bison *Bison bison* or caribou *Rangifer tarandus* show low network modularity. These species are likely to fit with the outcomes found in the ‘baseline scenario’, i.e. with no age structure in between-group contacts.

**Table 2:**
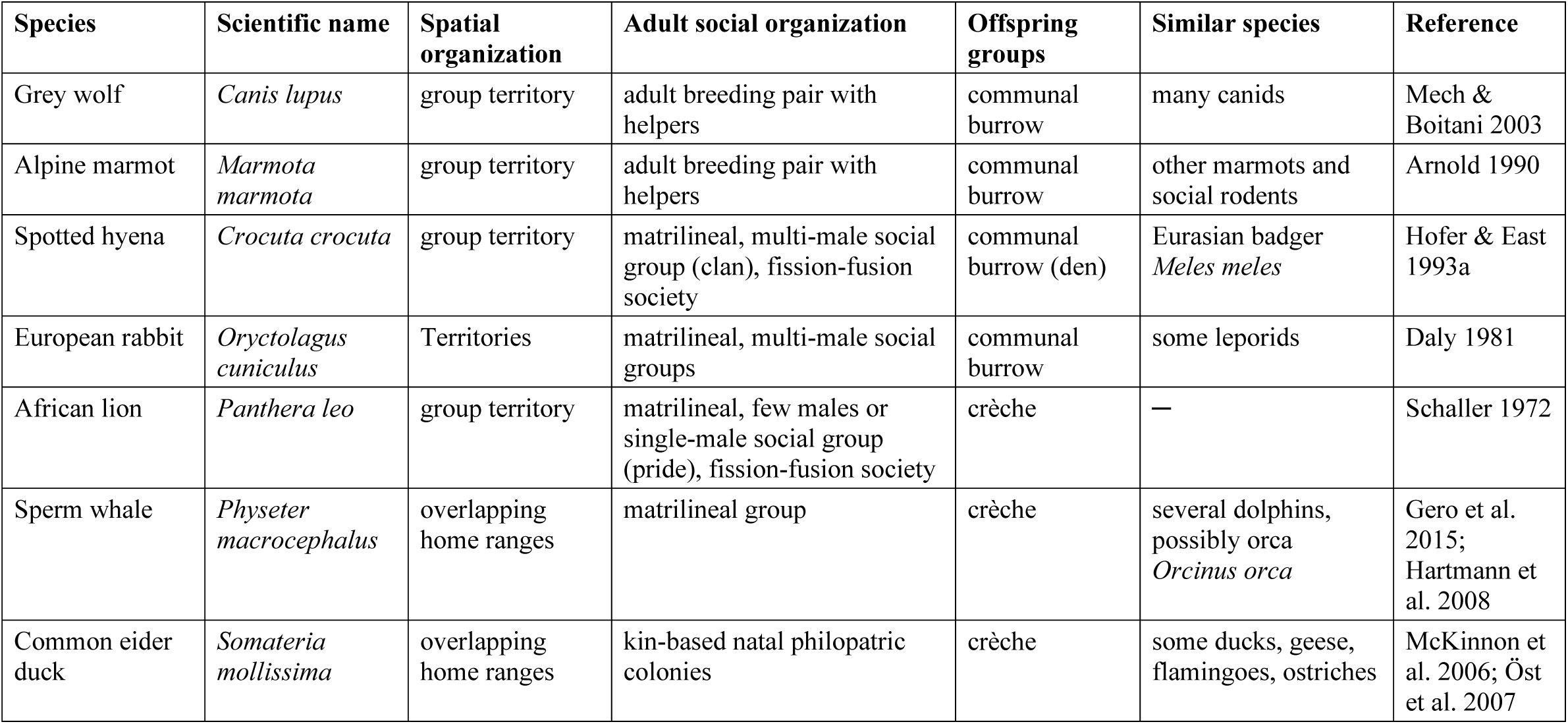
Examples of mammalian and avian species with social systems where distinct offspring units are looked after by adults in their own group and where the offspring have very little or no contact to members of other groups. These examples include species where offspring are stationary (i.e. within a den or burrow) or not (i.e. within a territory, but moving around within its boundaries).

In our model, individuals surviving pathogen infection acquired lifelong immunity. One typical example of such a pathogen in group-forming wildlife is canine distemper virus (CDV) (Appel and Summers 1995) or the now eradicated rinderpest (Daszak et al. 2000). Both morbiliviruses are highly contagious and some strains can cause massive mortality in wildlife. For example, a CDV epidemic in East Africa in 1993/1994 caused important declines of African lion and spotted hyena populations (Roelke-Parker 1996, Marescot et al. 2018, Benhaiem et al. 2018). CDV infects a broad range of mammals, mostly carnivores (Beineke, Baumgärtner, & Wohlsein, 2015, Deem, Spelman, Yates, & Montali, 2000). Even though many viruses are expected to induce lifelong immunity in their hosts, immunity can actually wane over time or provide only incomplete protection against reinfection (Morris et al. 2015). This is often difficult to verify in wild, free-ranging populations, as it requires repeated serological samples to measure antibody levels even in old animals. Even so, we expect our results to hold if immunity against the pathogen is maintained for several years. The impact of the length of the protective period of immunization (lifelong or less) could also be tested specifically with our model.

To conclude, our results show that the prolonged care of juveniles in communal nurseries or similar structures can reduce disease persistence in host populations of group-living species. A further development of the model would involve assessing the influence of hosts producing young continuously throughout the year (e.g. spotted hyenas) vs seasonal breeders (e.g. wild boars) on disease persistence. Birth patterns and the influx of new susceptible juveniles are indeed key drivers of infection with pathogens inducing lifelong immunity (Glass et al. 2011, Kramer-Schadt et al. 2009). For seasonal breeders, susceptible juveniles are only ‘available’ for infection for a few weeks or months, which might accelerate epidemic fade-out in comparison to species reproducing continuously.

## Supporting information

Supplementary Information-Keeping the kids at home

## DATA ACCESSIBILITY

This article has no additional data. The model script available on GitHub: https://github.com/LucileMarescot/RestrictingOffspringBetweenGroupContact_IBM.

## AUTHOR’S CONTRIBUTIONS

LM, MF, SKS and MLE designed the study. LM and MF developed the IBM, and carried out the statistical analyses and drafted the manuscript with SKS. SB contributed critically to data interpretation, revised the manuscript extensively and prepared the manuscript for submission. MLE and HH critically revised the manuscript. All authors gave final approval for publication and agree to be held accountable for the work performed therein.

## ACKNOWLEDGEMENTS

We are very thankful to Cédric Scherer for producing the figures, to Sonja Metzger for formatting the references and to Sylvain Gandon for his advice on the model conceptualization.

