## Supplementary Information-Keeping the kids at home for "‘Keeping the kids at home’ can limit the persistence of contagious pathogens in social animals"

[*] equal contribution

[‡] equal contribution

This document presents our procedure to ensure similar values of the basic reproduction number ($R_0$) between the host population with age-dependent between-group contact (i.e. 'ORC [offspring with restricted between group contact] scenario') and that with age-independent between group contact (i.e 'baseline scenario').

To compare the outcomes between the two types of host populations considering all parameter combinations, we controlled for similar values of $R_0$. Prior to any simulation, we calculated in the baseline scenario, the $R_0$ given the combination of pathogen traits and host traits. We then adjusted the infectivity of the pathogen in the ORC scenario to reach the same $R_0$ value as in the baseline scenario.

To do so, we first calculated the effective infection period $IL_{eff}$ for a given infected individual host. This can be expressed as the weighted average probability of dying over the maximum infection length added to the probability that the individual is still alive at the end of the infection length and has recovered:

$$IL_{eff} = \sum_{j=0}^{IL-1} j * (1-(1-\alpha)\phi) * [\phi(1-\alpha)]^{j} + IL * [\phi(1-\alpha)]^{IL} \qquad \text{(S1)}$$

We then determined the stable proportion of juveniles $P_{juv}$ in a population at carrying capacity by obtaining the geometric progression of the weekly probability of dying $(1 - (1 - \phi)$ before reaching the age at first between-group contact *age-contact*.

$$P_{juv} = \sum_{i=1}^{age-contact-1} (1-\phi) * (\phi)^{i-1} \qquad \text{(S2)}$$

We can thus express $R_0$ as the expected number of individuals within or between groups that are infected by the first infected individual during the maximum infection length. Assuming frequency dependent transmission and assuming that the first infected individual is an adult, $R_0$ can be determined as the expected number of individuals (juveniles and adults) that become infected during the effective infection length. This number is itself determined by the proportion of juveniles within a group and the probability that individuals belonging to the same group as the first infected individual to become also infected during its effective infection length (see (S1)). In addition, we must consider the expected number of adults that become infected during the effective infection length, which depends on the proportion of adults the probability that each of them become infected by the first infected individual and the pathogen infectivity *inf*.

$$R_0^{adult} = IL_{eff}\left[\left(1-\frac{1}{exp\left(\frac{inf}{(Group.size-1)*P_{juv}}\right)}\right)(Group.size-1)P_{juv}+\left(1-\frac{1}{exp\left(\frac{inf.\pi}{K-(Group.size)*(1-P_{juv})}\right)}\right)\left(K-(Group.size)*(1-P_{juv})\right)\right]$$

(S3)

Assuming frequency dependent transmission and assuming that the first infected individual is now a juvenile, the $R_0^{juv}$ is then expressed as:

$$R_0^{juv} = IL_{eff}\left[\left(1-\frac{1}{exp\left(\frac{inf}{Group.size-1}\right)}\right)(Group.size-1)\right] \quad (S4)$$

Therefore the total $R_0$ can be written as the sum of the $R_0$ of infected adults given that the first infected individual is an adult (red part of eqution S5) and the $R_0$ of juveniles given that the first infected individual was a juvenile (red part of equation (S5)):

$$R_0 = R_0^{juv} * P_{juv} + R_0^{adut} * (1-P_{juv}) \quad (S5)$$

Assuming that we approximate the real probability of within and between group infections (blue part of equation S3 and S4) by the expected number of individuals that become infected within and between the groups (blue part of equation S6) we may simplify the equation as follows:

$$R_0 = \mathrm{IL}_{eff}.\left[\frac{inf}{(Group.size-1)}\right](Group.size-1) * P_{juv} + \mathrm{IL}_{eff}.\left[\left[\frac{inf}{(Group.size-1)*P_{juv}}\right](Group.size-1) * P_{juv} + \left[\frac{inf.\pi}{K-(Group.size-1)*(1-P_{juv})}\right](K - (Group.size-1) * (1-P_{juv}))\right] * (1-P_{juv})$$

After simplification, this is equivalent to :

$$R_0 = \mathrm{IL}_{eff}.inf * P_{juv} + \mathrm{IL}_{eff}.[inf + inf.\pi] * (1-P_{juv})$$

Which is also equivalent to:

$$R_0 = \mathrm{IL}_{eff}.inf.P_{juv} + \mathrm{IL}_{eff} inf + \mathrm{IL}_{eff} inf.\pi - \mathrm{IL}_{eff} inf P_{juv} - \mathrm{IL}_{eff} inf.\pi.P_{juv}$$

Finally, we obtained the following equation for an approximation of $R_0$

$$\mathrm{R}_0 = inf.\mathrm{IL}_{eff}[1 + (1 - P_{juv})\pi] \quad \text{(S6)}$$

Therefore, for any combination of host and pathogen parameter, infectivity *inf* can be determined and adjusted in order to control $R_0$ using the following equation :

$$inf = \frac{\mathrm{R}_0}{.\mathrm{IL}_{eff}[1+(1-P_{juv})\pi]} \quad \text{(S7)}$$
